# *CRYPTID-exon*: streamlined detection of cryptic exons from RNA-seq data

**DOI:** 10.64898/2025.12.17.694983

**Authors:** Eraj S. Khokhar, Kaitlyn Brokaw, Zachary J. Kartje, Marina Krykbaeva, Ezequiel Calvo-Roitberg, Adam K. Hedger, Jonathan Lee, Atish Wagh, Jonathan K. Watts, Athma A. Pai

**Author notes:** These authors contributed equally (Eraj S. Khokhar, Kaitlyn Brokaw). current address: Alexion Pharmaceuticals (AstraZeneca Rare Disease), Cambridge, MA. current address: Korro Bio, Cambridge, MA.

## Abstract

Cryptic splicing has emerged as a pervasive feature of mammalian gene expression, with recent studies uncovering thousands of previously unannotated splice sites. Despite its prevalence, the functional consequences of this hidden layer of splicing remain largely unknown due to challenges in identifying the exact exonic regions introduced into mRNA transcripts. Here, we introduce a novel computational approach, *CRYPTID-exon*, that accurately predicts exon boundaries by modeling RNA-seq read coverage around empirically derived splice sites. We use *CRYPTID-exon* to identify and characterize thousands of cryptic exons in nascent and mature RNA from human cells. Additionally, we demonstrate that *CRYPTID-exon* is well powered to identify exons that are sensitive to translation-mediated degradation processes. Finally, given the growing interest in leveraging cryptic exons to modulate gene expression levels, we use our approach to identify cryptic exons in disease-relevant genes. We see that targeting these cryptic exons with splice-switching antisense oligonucleotides (ASOs) can alter gene expression and splicing patterns of the parent genes. Our study provides a framework to systematically identify and characterize cryptic exons, which will enable downstream insights into their impact on mRNA stability and translation.

## INTRODUCTION

RNA splicing is a ubiquitous feature of eukaryotic genomes that allows a single gene to encode multiple RNA and protein isoforms, with differences in both regulation and function. While splicing was historically thought to be heavily regulated to ensure accurate mRNA and protein compositions, recent studies have described pervasive cryptic splicing in mammalian cells.^1–4^ We and others have observed that most genes have low-frequency cryptic splicing events that result in transcripts with cryptic exons.^1,2^ Inclusion of these cryptic exons can change RNA structure, perturb regulatory motifs, alter open reading frames, and/or introduce premature termination codons (PTCs).^1,2,5^ Together, these changes can impact RNA stability, nuclear export, and transcriptional output. The majority of cryptic isoforms are likely subject to rapid degradation, either through nuclear surveillance pathways or cytoplasmic quality-control mechanisms such as nonsense-mediated decay (NMD).^1,2^ Crucially, the functional consequences of cryptic splicing events heavily depend on the sequence content of the newly included exonic regions. Systematic identification and characterization of cryptic exons is thus necessary for understanding how they shape gene expression and splicing profiles.

Cryptic exons have emerged as important disease drivers and thus therapeutic targets. For example, the RNA-binding protein TDP-43 often binds to pre-mRNA to suppress cryptic splicing.^6,7^ TDP-43 mis-localization, as is commonly seen in ALS and some forms of Alzheimer’s disease, leads to cryptic exon inclusion—including in Stathmin 2^8,9^ and UNC13A^10–12^—that causes pathologically important loss of functional protein.^13–15^ Similarly, repeat expansions in *FMR1*^16^ and *HTT*^17^ have also been associated with adverse pseudoexons that interfere with healthy gene expression and likely contribute to disease pathology in Fragile X syndrome and Huntington’s disease, respectively.

There has also been recent interest in profiling cryptic exons to redirect splicing for therapeutic interventions. Antisense oligonucleotides (ASOs) that modulate splicing patterns have been demonstrated in the literature as well as the clinic to be viable strategies for the partial correction and rescue of various target genes.^18,19^ This involves targeting and skipping exons whose inclusion leads to transcript degradation or altered reading frames. For instance, an ASO customized for a single Batten disease patient was able to reduce inclusion of a PTC-containing cryptic exon in *CLN7* and reduce seizure frequency by 80%.^20^ Similarly, four FDA-approved ASOs targeting exons within the dystrophin gene drive skipping of exons to restore reading frame disruptions created by disease-driving mutations.^21^ Finally, most genes likely have significant background expression of nonproductive, cryptic-exon-containing transcripts that can be targeted to increase expression of productive isoforms and thus increase overall gene expression. This strategy can be used to compensate for a loss-of-function disease-relevant mutation in the other allele.^22,23^ Together, these approaches hold great therapeutic promise but are all reliant on robustly identifying cryptic exons across many cellular and disease contexts.

Despite significant relevance for both biological and therapeutic purposes, identifying cryptic exons remains technically challenging. They are typically found at low abundance, making it difficult to delineate the full composition of a cryptic exon using PCR or high-throughput sequencing.^24,25^ Since cryptic isoforms are unstable and rare in mature RNA, recent work has turned to identifying splicing intermediates in nascent RNA before they are degraded.^1,2^ However, the presence of substantial unspliced molecules in nascent RNA further confounds the ability to easily identify cryptic exon boundaries. Several strategies have been employed to overcome these obstacles in mature and nascent RNA, including (1) manual inspection of read coverage,^6,20,26^ (2) identification of likely PTC-containing isoforms using sequence context^22^ or experimental enrichment,^26^ and (3) the depletion of regulatory factors (e.g. TDP-43,^6^ hnRNP L,^27^ PTBP1/2^28^) known to suppress cryptic splice site usage. This final experimental approach is often followed by constructing experiment-specific transcriptomes,^6,27^ and/or quantifying differential exon usage^10,29^ to identify cryptic exons. More broadly, methods for exon identification often rely on similarity to known coding sequences,^30,31^ homology-based approaches,^32^ and/or *ab initio* prediction of gene features.^33^ Together these approaches are labor-intensive, insufficiently powered to capture low-frequency events, dependent on cellular perturbations, or biased towards identifying cryptic exons with specific functional consequences.

We aimed to address this unmet need in transcriptomic analyses by developing a computational method to systematically identify cryptic exons using standard high-throughput sequencing datasets. We modeled read coverage around empirically derived splice sites to enable the detection of low-frequency cryptic exons from short-read RNA sequencing, including in nascent RNA. Using this approach, we identified thousands of cryptic exons, characterized their genomic properties, and tested the ability to target cryptic exons with splice-switching ASOs. Our work provides a framework for high-throughput cryptic exon profiling and highlights the need to further understand this underappreciated feature of gene expression.

## RESULTS

Despite advances in high-throughput sequencing of RNA that have uncovered pervasive cryptic splice site usage, the identification of exons that result from these sites remains a challenge for three main reasons. First, short RNA-seq exon junction reads that delineate spliced molecules generally span only one exon boundary, making it difficult to discern the full sequence composition of an exon. Second, since cryptic exons tend to be present at low relative levels in final transcripts, it is hard to distinguish exonic signal from noise in intronic regions. And finally, despite the promise of full-length long-read sequencing, the low coverage and high error rates inherent to these approaches complicate the identification of the precise boundaries for lowly expressed cryptic exons.^25,34^

To overcome these challenges, we developed a computational framework, CRYPTic IDentification of exons (*CRYPTID-exon)*, to predict the boundaries of cryptic exons using short-read RNAseq data. This framework is built around two critical features. First, *CRYPTID-exon* uses empirically derived splice sites as a starting point for cryptic exon identification (**Fig. 1A**). Second, anchoring on these sites, we use a probabilistic changepoint algorithm^35,36^ to identify a region with significantly higher read coverage than coverage in flanking intronic regions (Methods). This strategy relies on the premise that the 3’ and 5’ boundaries of a bona fide cryptic exon will show steep increases and decreases in coverage, respectively, relative to surrounding intronic regions. These will be identified as changepoints in the distribution of local read coverage.

**Figure 1.**
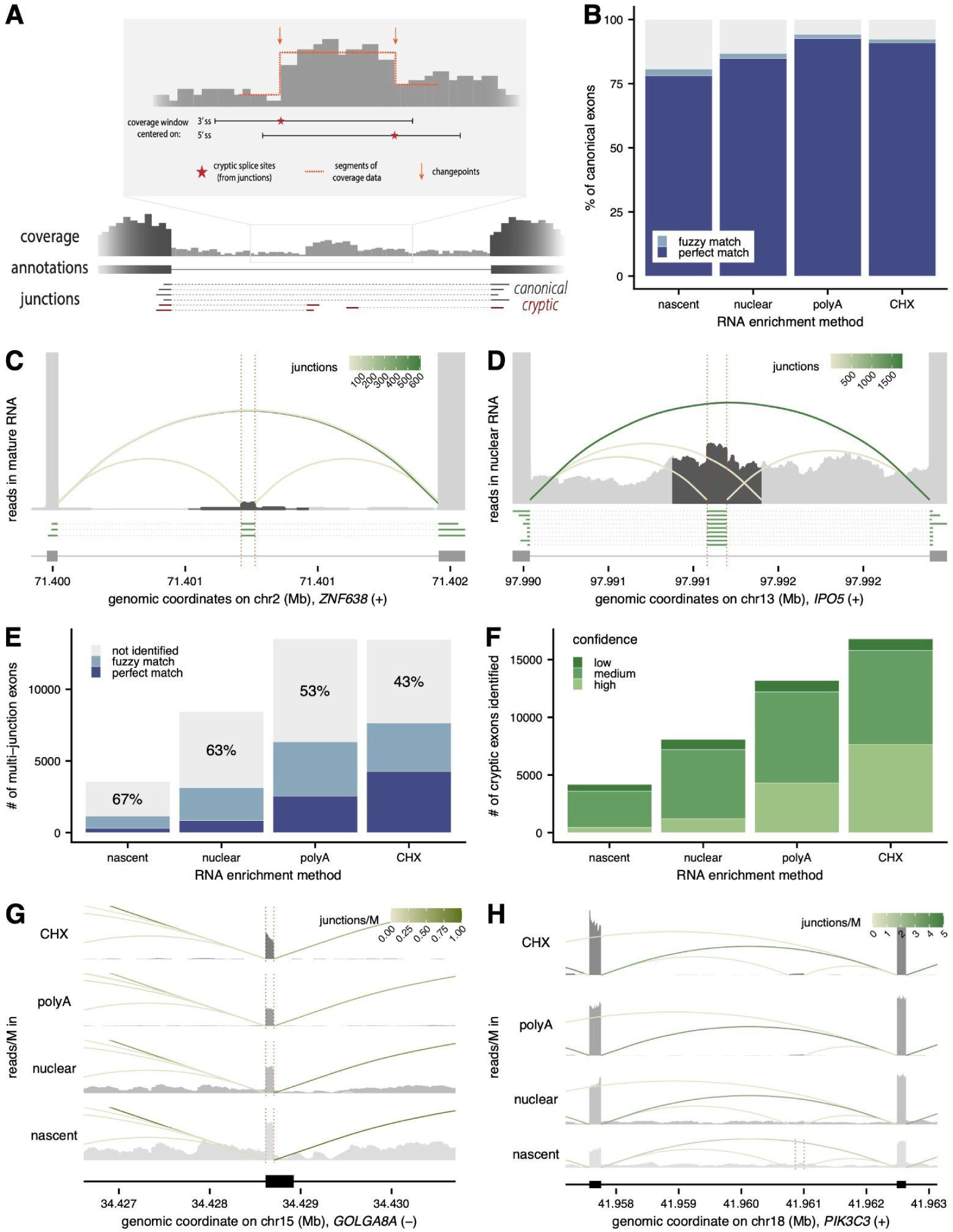
*CRYPTID-exon* identifies cryptic exon boundaries using short read RNA-seq data. **(A)** Schematic of the *CRYPTID-exon* framework, where RNA-seq coverage in a window centered around empirical splice sites is used to identify changepoints. **(B)** Percentage of canonical exons correctly identified by *CRYPTID-exon*, with either perfect (*dark blue*) or fuzzy boundary (*light blue*) matches, across RNA enrichment methods. Canonical exons are expressed in mature RNA in K562 cells and ≤200 nt. **(C, D)** Representative examples of cryptic exons identified in *ZNF638* from mature RNA **(C)** and *IPO5* from nuclear RNA **(D)**. RNA-seq read coverage (*top*) for a wider window (*light grey*) and the specific window used for running *CRYPTID-exon* (*dark grey*), with curved lines representing exon-exon junction reads (colored by abundance). Green solid and dotted lines (*middle*) show multi-junction reads and gene annotations are shown at the bottom. **(E)** Percentage of cryptic exons from multi-junction reads correctly identified by *CRYPTID-exon*, with either perfect (*dark blue*) or fuzzy boundary (*light blue*) matches. **(F)** Numbers of cryptic exons identified by *CRYPTID-exon* in K562 cells across RNA enrichment methods, separated by low (*black*), medium (*green*), and high (*light green*) confidence categories. **(G, H)** Cryptic exons identified in *GOLGA8A* **(G)** and *PIK3C3* **(H)** in K562 cells. Read coverage (*top, grey*) and junction reads (*curved lines*, shaded by abundance) are shown across RNA enrichment methods, with dotted lines representing datasets in which the cryptic exon was identified. Gene annotations are shown at the bottom.

### *CRYPTID-exon* can accurately and precisely predict exon boundaries

The main empirical parameters that underlie the *CRYPTID-exon* framework to identify exons from short-read RNA-seq data are the size of the coverage window within which changepoints are identified and the relative coverage of the exon compared to flanking regions. We used canonical exons expressed in K562 erythroleukemia cells (Methods) as a gold-standard set of exons to calibrate these parameters. Using canonical exons of different lengths, we first assessed the necessary size of a coverage window around a splice site to confidently identify changepoints (**Fig. 1A**). For both 3’ and 5’ss centered windows, we see that a window extending 100nt into the putative intronic region and 200nt into the putative exonic region performs the best for predicting exons that are 200nt or less (**Supplementary Fig. 1A**). Using this window, we are able to precisely identify the boundaries for 93% of randomly sampled canonical exons that are 200nt or less and an additional 1.7% with “fuzzy” boundaries (≤ 20nt from the precise boundary; **Supplementary Fig. 1B**). We further applied *CRYPTID-exon* to canonical exons identified from RNA enrichment methods that sample RNA across cellular compartments: nascent RNA from the nucleus, nuclear RNA, mature RNA after polyA selection, and cycloheximide-treated RNA (CHX) to examine the effects of translation-dependent degradation mechanisms like nonsense mediated decay (NMD).^2^ In these data, we identify boundaries for 78-93% of canonical exons that are 200nt or less (81-94% conditioning on a perfect boundary match; **Fig. 1B**). We identify the most exons in mature RNA (+/- CHX), as expected given the increased intronic read coverage contributed by unspliced transcripts in nascent and nuclear data.

Next, we wanted to assess our sensitivity to detect cryptic exons. To do so, we used cryptic exons identified from the limited number of multi-junction reads that span more than one splice junction (at least one of which includes a cryptic splice site), thereby precisely delineating the exonic boundaries. For instance, a cryptic exon in intron 15 of *ZNF638* is supported by three multi-junction reads and increased read coverage in mature RNA (**Fig. 1C**), while a cryptic exon in intron 9 of *IPO5* is supported by nine multi-junction reads and moderately increased read coverage despite substantial contributions from unspliced transcripts in nuclear RNA (**Fig. 1D**). For both of these examples, *CRYPTID-exon* identifies cryptic exon boundaries that match the boundaries from multi-junction reads. We see that *CRYPTID-exon* again performs best for these exons using a 300nt window around the splice site, extending 100nt into the intron and 200nt into the putative exon, regardless of the RNA enrichment method from which they are identified (**Supplementary Fig. 1C**). However, *CRYPTID-exon* is only able to identify 33 - 57% of all multi-junction read delineated cryptic exons with some confidence (**Fig. 1E**), likely because the majority of these exons are still expressed quite lowly. Consistently, we see better identification for multi-junction exons with higher multi-junction read support and thus increased relative cryptic exon usage (**Supplementary Fig. 1D**).

### *CRYPTID-exon* identifies exons associated with thousands of cryptic splice sites

To identify exons associated with cryptic sites genome-wide, we used *CRYPTID-exon* to scan for exons anchored on every 3’ and 5’ss identified across RNA enrichment methods in K562 cells. We identified splice sites using *CRYPTID-SS*, a computational framework we previously developed to identify high confidence canonical and cryptic splice sites in high-throughput sequencing data^2^. After collapsing exons predicted from 3’ and 5’ss (Methods), we identified 25,619 unique exons associated with cryptic sites across RNA fractions (**Fig. 1F, Supplementary Table 1**), with the most exons identified in mature RNA fractions. Of note, we identify ∼30% greater cryptic-site-associated exons in CHX-treated RNA, suggesting that *CRYPTID-exon* is well powered to detect NMD-sensitive cryptic exons. As expected given the relative usage of canonical vs. cryptic sites, *CRYPTID-exon* identifies exons associated with 60-65% of canonical 3’ss but only 5-27% of cryptic 3’ss identified across RNA fractions (**Supplementary Fig. 1E**). For most cryptic site-associated exons, predicted exon boundaries are within 10nt of the associated splice site (**Supplementary Fig. 1F**), with a slightly larger distribution around 3’ss that might reflect the usage of NAGNAGs.^37^

To confirm the boundaries of cryptic exons predicted by *CRYPTID-exon*, we selected two exons for independent validation. First, we chose a cryptic exon that uses a cryptic alternative 5’ss that modifies exon 2 of *GOLGA8A*, which was identified and can be seen to be expressed across all RNA fractions in K562 cells (**Fig. 1G**). Using amplicon sequencing with primers designed within the cryptic exon and flanking exons (Methods; **Supplementary Table 2**), we confirmed the boundaries of this cryptic exon (**Supplementary Fig. 2A**). Second, we looked at a novel cryptic exon within intron 2 of *PIK3C3*, which was identified by *CRYPTID-exon* in nascent RNA data from K562 cells but can also be visually seen in mature RNA (**Fig. 1H**). Amplicon sequencing again confirmed the boundaries of this cryptic exon (**Supplementary Fig. 2B**).

Since *CRYPTID-exon* allows for some uncertainty in splice site and exon boundary concordance, we categorized the identified exons into three confidence levels (**Supplementary Fig. 2C**): (1) *high confidence exons*, where exon boundaries and splice sites perfectly match between exons identified independently from 3’ and 5’ss, (2) *medium confidence exons*, where either exon boundaries or splice sites perfectly match between 3’ and 5’ss-identified exons, and (3) *low confidence exons*, where neither are perfect matches but all metrics are within 20nt (Methods). On average, low confidence exons have the lowest number of supporting junction reads for both 3’ and 5’ss (**Supplementary Fig. 2D**) and the lowest relative exonic coverage relative to flanking intronic reads (**Supplementary Fig. 2E**). Thus, these exons are likely those used at lowest frequencies. To understand characteristics of cryptic site-associated exons, we used only medium and high confidence exons for our downstream analyses, but believe that low confidence cryptic exons are good starting points for experimental confirmation in genes of interest.

### Novel cryptic exons are shorter and less conserved than annotated exons

When identifying exons anchored by cryptic splice sites, it is still possible to identify known or partially known exonic regions. Thus, we first compared the exons associated with cryptic sites with the boundaries of reference exons (Methods) and identified four overlap categories (**Fig. 2A**): (1) *annotated exons*, where a cryptic site-associated exon exactly matches (within 2nt) a reference exon, (2) *partial exons*, where the boundaries of a cryptic site-associated exon fall within the boundaries of a reference exon, (3) *extended exons*, where the boundaries of a cryptic site-associated exon overlap a reference exon but add a novel region onto either the 5’ or 3’ end, and (4) *novel exons*, where the boundaries of a cryptic site-associated exon do not overlap a reference exon. We find that, across RNA enrichment methods, 79-83% of *CRYPTID-exon* predictions associated with only one cryptic site are categorized as annotated exons (**Fig. 2B**), with the majority of these being associated with high-fidelity cryptic 3’ss paired with canonical 5’ss (**Supplementary Fig. 3A**) that are found within 10nt of the annotated splice site (**Supplementary Fig. 3B**). This suggests that the majority of exons associated with only one cryptic site might arise from pervasive NAGNAG usage at very low frequency^37^ and *CRYPTID-exon* is not sensitive enough to differentiate between exons deriving from these closely positioned alternative 3’ or 5’ss.

**Figure 2.**
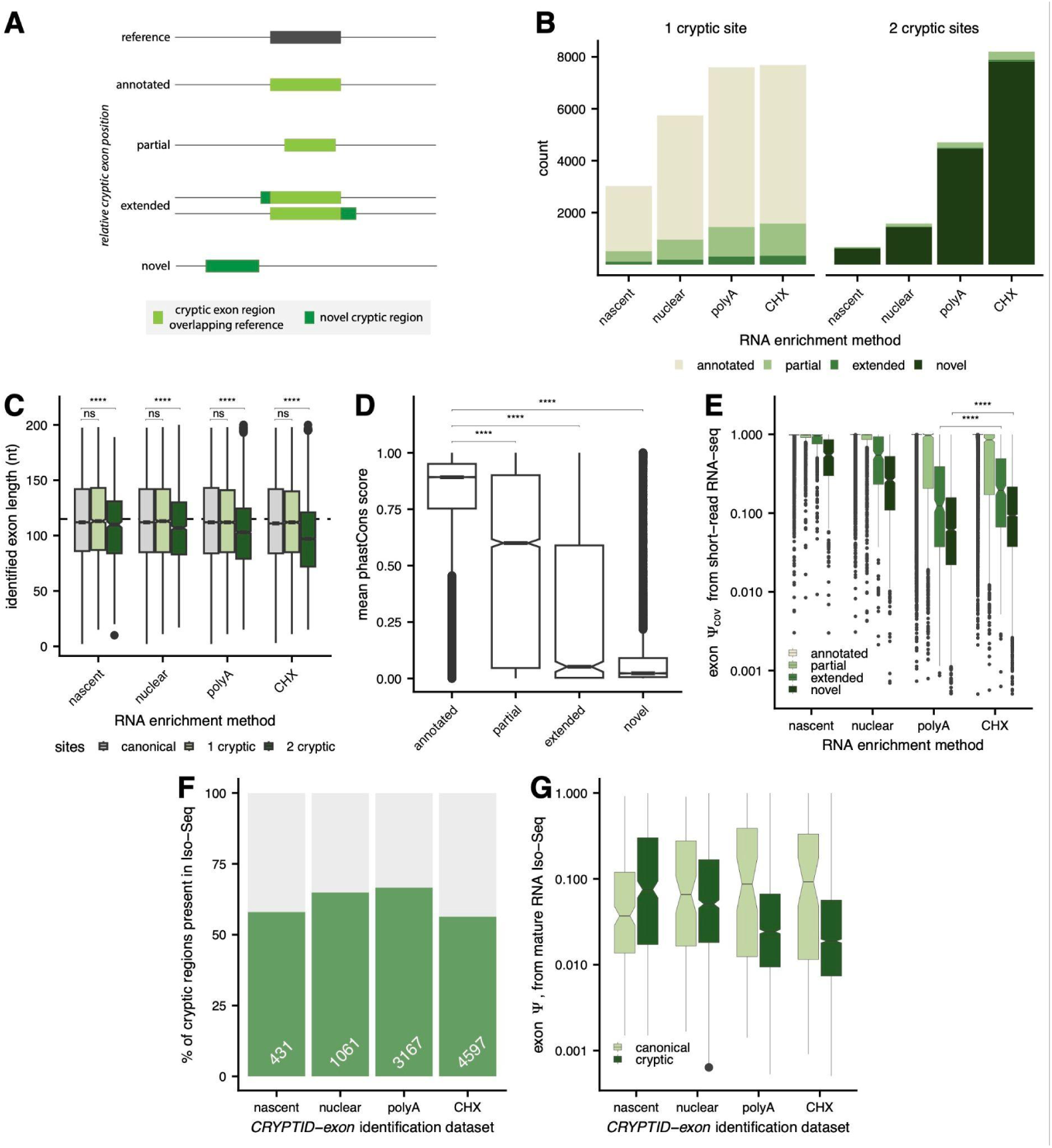
***CRYPTID-exon* identifies novel exonic regions associated with cryptic splice sites. (A)** Schematic of the overlap between identified exons and annotated exons, where cryptic exons are categorized based on full, partial, or no overlap. **(B)** Number of cryptic exons with either 1 (*left*) or 2 cryptic splice sites (*right*) assigned to each overlap category across RNA enrichment methods. **(C)** Distribution of exon lengths identified by *CRYPTID-exon* for exons with canonical splice sites (*grey*) or cryptic sites (*shades of green*) across RNA enrichment methods. **(D)** Distribution of mean phastCons scores for cryptic exons across overlap categories. **(E)** Distribution of exon Ψ_cov_ values for cryptic exons across overlap categories and RNA enrichment methods. **(F)** Percentage of novel cryptic regions that are supported by read coverage in long-read Iso-Seq data. **(G)** Distribution of exon Ψ values estimated from mature RNA Iso-Seq data from K562 cells for novel canonical (*light green*) or cryptic (*dark green*) exons identified in different RNA enrichment methods. Significance was assessed with a Mann-Whitney U Test. **** adjusted p-value < 0.0001.

In contrast, 91-95% of predictions associated with two cryptic sites are novel exons (**Fig. 2B**), which tend to use two high-fidelity cryptic splice sites (**Supplementary Fig. 3A**). These novel exonic regions tend to be significantly shorter than annotated exons associated with canonical sites (**Fig. 2C**, **Supplementary Fig. 3C**). Notably, the median length of novel exons found in nascent and nuclear RNA is 8nt longer than novel exons found in mature RNA, while the median length of novel exons in CHX-treated RNA is 6nt shorter. This is consistent with previous observations that shorter exons with premature stop codons are more likely to be sensitive to NMD.^38^ In contrast, novel extended regions added to reference exons by alternative 3’ and 5’ splice sites are longer in CHX-treated RNA (**Supplementary Fig. 3D**). Finally, we see that partial, extended, and novel cryptic exons are all significantly less conserved than annotated exons (**Fig. 2D**). Novel exons exhibit conservation signatures that are the same as flanking intronic regions (**Supplementary Fig. 3E**), suggesting that these regions are less likely to encode for functional RNA regions.

### Cryptic exons are used infrequently in mature RNA

We next wanted to quantify how often cryptic exons were included in spliced RNA transcripts. To do so, we developed a “percent spliced in” metric based on read coverage (Ψ_cov_) that accounts for background unspliced transcripts using flanking intronic read coverage (Methods). Using this metric, we see that extended and novel cryptic exonic regions are used relatively infrequently (**Fig. 2E**), especially in mature RNA where they are only included in ∼10% of transcripts on average. In CHX-treated RNA, the inclusion levels for both extended and novel cryptic exons significantly increase by approximately 60%, supporting the idea that a fraction of these exons trigger translation-dependent degradation processes (i.e. NMD) upon their usage. The use of an alternative metric that relies solely on junction reads supporting inclusion or exclusion of cryptic exons (Ψ_junc_, Methods) shows similar extents of cryptic exon usage (**Supplementary Fig. 3F**).

An orthogonal and more direct high-throughput method to quantify exon inclusion is long-read sequencing (LRS), which allows for placement of an exonic region in the full isoform context. We used publicly available Iso-Seq LRS data from K562 cells^39^ and first assessed how often we observe *CRYPTID-exon* identified exons in these data. We see that, after filtering for expressed genes and complete reads (Methods), 46-66% of novel exons identified across RNA enrichment methods are also present in LRS reads (**Fig. 2F**). For instance, there are 60 and 3 reads that contain the cryptic exons that we identified in both *GOLGA8A* and *PIK3C3*, respectively (**Supplementary Fig. 3G-H**; 23% and 1% of total reads for the two genes). More broadly, we see that novel regions that are associated with either canonical or cryptic splice sites are included in less than 10% of mature transcripts, on average, and that regions identified in mature RNA (+/- CHX) are the least included (**Fig. 2G**). Notably, given the limitations of long-read sequencing data, we identify an order of magnitude fewer novel regions in the LRS data, supporting the continued need for high coverage short-read data to comprehensively identify cryptic exons.

### Novel cryptic exons are likely to introduce premature stop codons

To understand the potential functional consequences of cryptic exon usage, we looked at the positioning of cryptic exons within genes. While non-annotated cryptic exons are enriched within non-coding transcripts and untranslated regions (UTRs), the majority of cryptic exons are associated with coding regions (either modifying coding exons or within introns found in open reading frames; **Fig. 3A**, **Supplementary Fig. 4A**). Thus, cryptic exons may often change or disrupt translational reading frames. We first asked if they are likely to disrupt reading frames by evaluating whether inclusion of the cryptic exon is likely to change codon usage within the reading frame. We see that, as expected, annotated exons within coding regions are more likely to be evenly divisible by three (**Fig. 3B**, **Supplementary Fig. 4B**), suggesting that differential usage of these exons is less likely to shift translation frame or codon identity. However, non-annotated cryptic exons do not show the same trend, suggesting they might be more likely to have frame-shifting potential. To directly test if these exons introduce premature stop codons, we assigned each cryptic exon within a coding region a reading frame based on the nearest upstream coding exon and scanned for the presence of known stop codons (Methods). We see that novel cryptic exons are enriched for stop codons relative to canonical exons (**Fig. 3C**), suggesting that these exons are likely to introduce premature stop codons that can trigger NMD.

**Figure 3.**
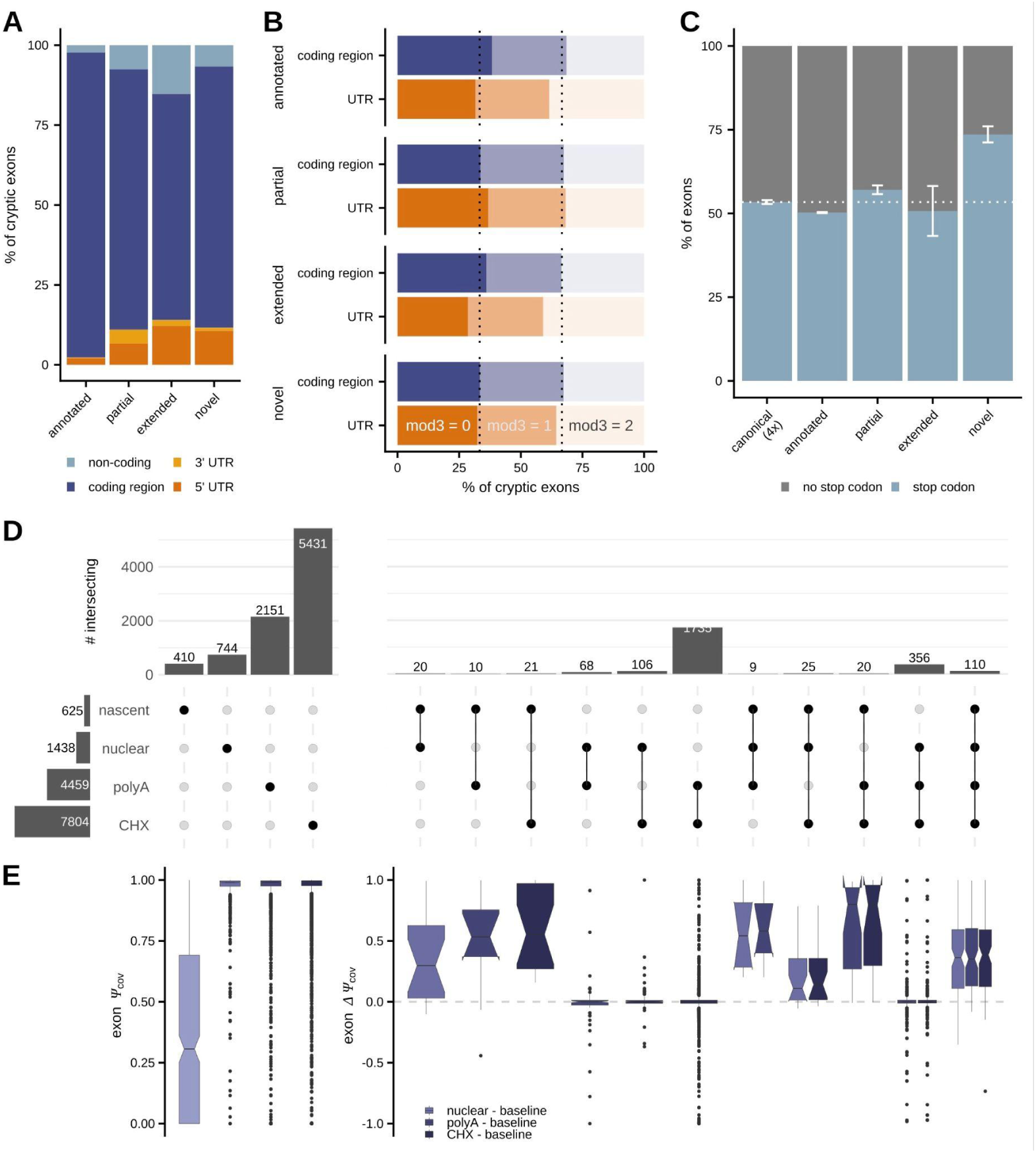
Novel cryptic exons likely trigger nonsense-mediated decay. **(A)** Percentage of cryptic exons that are found in non-coding genes (*light blue*), coding regions (*dark blue*), 3’UTRs (*light orange*), or 5’ UTRs (*dark orange*) across overlap categories. **(B)** Percentage of cryptic exons in coding regions (*blue*) or UTRs (*orange*) whose length is evenly divisible by 3 (mod3 = 0; maintaining frame; *darkest shade*) or not evenly divisible by 3 (mod3 = 1 or mod3 = 2; *lighter shades*) across overlap categories. **(C)** Percentage of cryptic exons predicted to have in-frame stop codons (*blue*) across overlap categories, with error bars reflecting variation across exons identified in different RNA enrichment methods. As a control, the same percentage was calculated in four samples of canonical exons (*left*). **(D)** The number of novel cryptic exons detected in only one (*left*) or multiple (*right*) RNA enrichment method, with the total number of novel cryptic exons within each RNA enrichment method shown as bars on the left. **(E)** Distribution of exon Ψ_cov_ values (*left*) or exon ΔΨ_cov_ values (*right*) for novel cryptic exons that show different overlap patterns (as shown in D). Exon ΔΨ_cov_ values are estimated relative to the earliest RNA fraction in which the exon is identified.

### Thousands of novel cryptic exons are sensitive to translation-dependent degradation mechanisms

Given the observation that many cryptic exons may trigger NMD, we wanted to understand how the usage of cryptic exons changed across the RNA lifecycle from nascent to translating mature RNA. For both novel and extended cryptic exons, we see that the majority of exons are only identified in one RNA enrichment method (**Fig. 3D**; **Supplementary Fig. 4C**) and these compartment-specific exons are included in the majority of transcripts (**Fig. 3E**; **Supplementary Fig. 4D**). Interestingly, we detect more than double the number of novel cryptic exons in CHX-treated RNA than in mature RNA, suggesting the usage of substantial numbers of NMD-sensitive cryptic exons. Mature RNA fractions (+/- CHX) share the most novel and extended cryptic exons – ∼1735 and 106 exons, respectively – and these exons are, on average, used at the same frequency in both fractions. These exons may have translational potential and are most likely stable cryptic exons with delayed splicing that is not yet observed in nascent or nuclear fractions. The next two largest groups of shared cryptic exons are those shared across nuclear and mature RNA fractions and all four fractions. For shared cryptic exons, the inclusion of the exon increases as the transcript gets more mature, as measured by the change in Ψ_cov_ (ΔΨ_cov_) relative to the earliest RNA fraction in which the exon was identified (**Fig. 3E**; **Supplementary Fig. 4D**).

Finally, we wanted to understand if cryptic exon usage was cell-type specific. To do so, we ran *CRYPTID-exon* on a matching dataset of RNA sampled across cellular compartments from KNS60 glioblastoma cells.^2^ (**Supplementary Table 1**) We see that greater than 60-66% of total canonical exons identified are shared across cell types and 71-82% of the remaining exons are were specifically identified in KNS60 cells (**Supplementary Fig. 4E**). In contrast, only 16-25% of cryptic exons are shared across cell types, with the highest amount of sharing seen in mature RNA and particularly CHX-treated RNA. This suggests that cryptic exon usage is likely to occur in a cell-type specific manner, either through stochastic or regulated splice site usage.

### ASO-mediated targeting of cryptic exons alters gene expression and splicing

The development of splice-switching ASOs has led to interest in identifying and targeting cryptic exons in disease-implicated genes to modulate gene expression levels. To evaluate the utility of our combined framework for identifying cryptic splice sites and exons within this context, we focused on cryptic splicing in genes known to cause haploinsufficiencies when one allele contains a mutation. Specifically, we chose 13 genes expressed in neuronal cells: *ENPP1*, *FOXP1*, *GRN*, *MAPT*, *MLLT3*, *NEK1*, *NF1*, *OPTN*, *PUM1*, *SYNGAP1*, *TBK1*, *TSC1*, and *TSC2*.

Since these genes are lowly to moderately expressed in KNS60 cells, we first enriched for target transcripts from nuclear RNA using probe-based cDNA pulldowns followed by high-throughput sequencing (**Fig. 4A**, **Supplementary Table 3;** Methods). The enriched samples show a 30 to 2,800-fold increase in abundance of the target genes relative to nuclear RNA (**Supplementary Fig. 5A**). Using these data, we ran *CRYPTID-SS* and *CRYPTID-exon* to identify cryptic splice sites and exons, respectively. The target genes have some of the highest number of junction reads supporting both cryptic and canonical splice sites (**Fig. 4B**; **Supplementary Fig. 5B**), providing confidence in the ability to robustly identify cryptic splicing patterns for these genes following targeted sequencing. We decided to further examine the three genes with the highest number of cryptic junction reads: *NF1*, *PUM1*, and *TBK1*. *CRYPTID-exon* identified 11 cryptic exons from *NF1,* 3 from *TBK1,* and 4 from *PUM1* with high or medium confidence (Methods). Focusing on the exons linked to the cryptic sites with the highest expression in these three genes (**Fig. 4C**), we validated the coordinates of these exons using amplicon sequencing as described above (**Supplementary Fig. 5C-E**).

**Figure 4.**
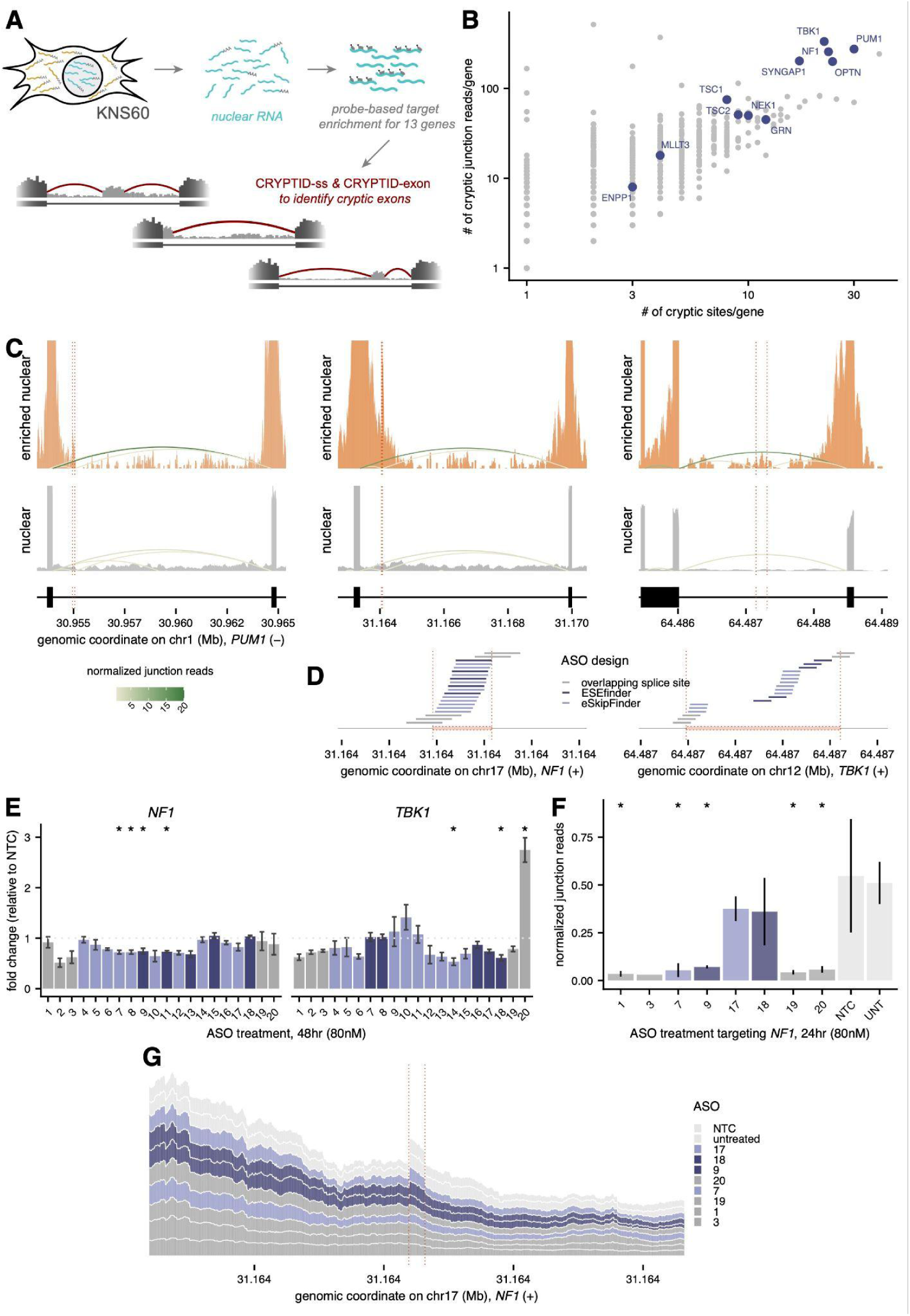
Gene expression levels and splicing patterns can be modulated by targeting cryptic exons with ASOs. **(A)** Schematic of probe-based enrichment strategy to identify cryptic sites and exons in disease-relevant genes. **(B)** The number of cryptic sites identified per gene (*x-axis*) vs. the number of cryptic junction reads identified per gene (*y-axis*) in enriched nuclear RNA, with target genes highlighted in blue. **(C)** Read coverage for *PUM1* (*left*), *NF1* (*middle*), and *TBK1* (*right*) in enriched nuclear (*top*, *orange*) and nuclear RNA (*bottom*, *grey*) samples, with curved lines representing exon-exon junction reads (colored by abundance). Dotted lines represent the boundaries of cryptic exons. **(D)** Positions of ASOs designed using ESEfinder (*dark blue*), eSkipFinder (*light blue*), or to directly overlap the cryptic splice sites (*grey*) for *NF1*(*left*) and *TBK1* (*right*). **(E)** Fold changes in gene expression levels estimated by RT-qPCR for *NF1* (*left*) and *TBK1* (*right*) relative to a non-treated control (NTC), following transfection with individual ASOs at 80nM for 48 hours. ASOs are numbered by their genomic position, with 1 and 20 representing the upstream-most and downstream-most ASOs, respectively. **(F)** Normalized junction reads associated with the 3’ss of the *NF1*-identified cryptic exon. (*y-axis*) in enriched nuclear RNA following ASO treatment or in control samples (*x-axis*). For G, significance was assessed with a T-test relative to the untreated sample. * FDR < 0.1. **(G)** Read coverage around NF1-identified cryptic exon (*y-axis*) in enriched nuclear RNA following ASO treatment or in control samples (*colors*).

Moving forward with the cryptic exons identified in *NF1* and *TBK1*, we used three different approaches to design splice-switching ASOs that tile the exon: (1) eSkip-Finder^40^ to identify ASO sequences with the highest predicted skipping efficacy within the exon, (2) ESEfinder^41^ to identify putative exon splicing enhancers to target with ASOs, and (3) manually designed ASOs designed to occlude predicted 5’ or 3’ss of the exon (**Fig. 4D, Supplementary Table 4**). We first evaluated whether treatment with individual ASOs resulted in gene expression changes in KNS60 cells. We see that all of the ASOs designed for *NF1* and the majority of ASOs for *TBK1* result in no change or slight down-regulation of gene expression for the target genes relative to non-targeting control ASO, regardless of ASO treatment duration (**Fig. 4E**; **Supplementary Fig. 6A, Supplementary Table 5**). Two ASOs for *TBK1*—one predicted to be efficacious by eSkip-Finder and another that overlaps the 5’ss—resulted in apparent up-regulation of gene expression, with the latter ASO (“ASO 20”) resulting in a 2.75 fold increase in mRNA abundance. However, this ASO was also associated with substantial decreased expression of the housekeeping gene, potentially contributing to its apparent *TBK1* activation (**Supplementary Fig. 6B**), as well as with changes in cellular phenotype suggesting some cytotoxicity. Off-target effects have previously been associated with other apparent gene activation observations^42^. To study whether pairs of ASOs would be more effective than single ASOs, and with a focus on ASOs that block the cryptic splice sites, we treated KNS60 cells with a mix of two ASOs —- one targeting the 3’ss and another targeting the 5’ss. However, the only combinations driving profound activation were pairs that included the potentially toxic ASO 20 (**Supplementary Fig. 6C,D**). As such, this small screen did not identify good hits for therapeutic gene activation of either target.

Nevertheless, we wanted to understand if targeting cryptic sites with splice-switching ASOs prevented splicing and reduced inclusion of the cryptic exon, even when gene activation was not observed. We chose 8 ASOs targeting the *NF1* cryptic exon, distributed across design approaches, and performed our target enriched sequencing strategy after ASO treatment (paired with untreated and non-targeting control samples). We see that treatment with five of the ASOs (but not the non-targeting control) significantly reduces splicing at the cryptic site (**Fig. 4F**) and we see a concordant decrease in the inclusion of the cryptic exon upon treatment with these ASOs (**Fig. 4G**; **Supplementary Fig. 6E**). Together, these results suggest that our combined framework can identify cryptic sites and exons that can be targeted with ASOs to alter gene expression and splicing patterns.

## DISCUSSION

Here, we describe *CRYPTID-exon*, a novel computational approach to identify and quantify genome-wide cryptic exon usage in mammalian cells. We validate our method using canonical exons, bona fide cryptic exons delineated by junction reads, and orthogonal sequencing approaches. Our results suggest that there are thousands of unannotated cryptic exons that are used at low levels across nascent and mature RNA populations. To our knowledge, this is the first systematic identification of novel exons across successive stages of RNA maturity, allowing for the profiling of previously unexplored mechanistic consequences of these cryptic exons in addition to any coding potential.

Despite the utility of *CRYPTID-exon*, we are still unable to identify exons associated with the majority of cryptic sites. This is likely because most cryptic sites are used at extremely low frequencies, resulting in exonic regions that appear in only a tiny fraction of transcripts and leading to an inability to confidently infer precise exon boundaries. Highly expressed cryptic splice site are also often alternatively paired with multiple splice site partners.^2^ This could create a range of possible exon boundaries that disperse the signal across many exon configurations and complicates the identification of a single, well-supported exonic region. While both of these issues affect the expression of cryptic exons, these exons are also likely being actively and rapidly degraded, leading to further insufficient coverage in either nascent or mature RNA populations to reconstruct the exon. Applying *CRYPTID-exon* to more datasets that enrich for cryptic splicing intermediates, through targeted sequencing and/or perturbation of degradation mechanisms, will be useful for exploring the range of as yet undiscovered cryptic exons.

The ability to identify cryptic exons using *CRYPTID-exon* is also constrained by exon length. The average vertebrate exon length is 137 nt^43^ but the average length of canonical exons that we are able to identify is 117nt. This is consistent with the limitations on the length of the coverage window used to scan for exons. We find that coverage windows extending into the exon for more than 200nt perform poorly, likely due the higher probability of overlapping alternative exon configurations and noisier background signal due to short reads. Longer high-throughput sequencing reads could in principle extend the range and complexity of exons that can be detected, particularly by increasing the abundance of multi-junction reads. Additionally, advances in dedicated long-read sequencing platforms would also aid cryptic exon identification. Current long read platforms still suffer from high error rates that complicate the precise identification of splice site boundaries. Existing pipelines to analyze long read data compensate for this by using known annotations or splice site sequences to correct or “realign” boundaries,^25,34^ but this limits their ability to discover new cryptic splice sites.^44^ Integrating information from high coverage short read and full-isoform long read datasets within our *CRYPTID-SS* and *CRYPTID-exon* frameworks might alleviate these issues.

We find that novel cryptic exons are more likely to contain stop codons. However, errors in predicted exon boundaries can obscure reading frame inference. It is also challenging to infer their reading frame and thus sequence-based sensitivity to NMD without knowing the full isoform context. Information about full length cryptic isoforms could also inform their impacts on transcript regulation and coding potential. Still, the observation that cryptic exon usage increases—in both number of exons detected and relative usage of each exon—during translation inhibition suggests that many of these isoforms are indeed subject to NMD, consistent with their low abundance in steady-state RNA and previous observations.^1,26^

Finally, our findings have implications for therapeutic modulation of cryptic splicing using ASOs. The identification of a targetable cryptic exon is a first step towards devising an ASO-based treatment strategy.^45^ Thus our *CRYPTID-exon* pipeline may be useful to use alongside guidelines and resources developed by others that drive variant identification and enable rare ASO drug development.^46–48^ Our preliminary observations indicate that ASOs targeting cryptic exons and their boundaries can drive skipping of the exons discovered using *CRYPTID-exon*. Whether this skipping of cryptic exons leads to up-regulation of the canonical isoform likely depends on the expression levels of the cryptic isoform and whether the cryptic isoform affects regulatory feedback loops. While evaluating the latter is beyond the scope of our pipeline, the application of *CRYPTID-exon* to identify and short-list highly included cryptic exons, especially in genes with pathological relevance, would be an effective strategy to identify candidate exons for further study. Our work thus lays the groundwork for developing predictive frameworks to interpret biological consequences of cryptic splicing, its role in post-transcriptional regulation, and the ability to modulate cryptic splicing as a targeted RNA-based therapy.

## METHODS

### Identification of cryptic exons with *CRYPTID-exon*

We developed a python-based framework to confidently identify the boundaries of cryptic exons using RNA-seq reads coverage. Briefly, this pipeline consists of the following five steps, each of which are detailed below: (1) obtain read coverage centered around empirically observed splice sites, (2) predict exon boundaries for each splice site using a changepoint detection algorithm, (3) combine cryptic information across 3’ and 5’ splice sites, (4) identify cryptic exons with low, medium, or high confidence. *CRYPTID-exon* takes as inputs a list of 3’ and 5’ splice sites and a read alignment file (.bam) and outputs a filtered table that contains predicted exons and associated splice sites.

#### Step 1: Read coverage in splice site windows

For each splice site in the user-provided list of splice sites, *CRYPTID-exon* gets read coverage in a window around the splice site from which to identify a putative cryptic exon associated with that splice site. Specifically, the pipeline uses bedtools coverage^49^ to get the number of reads, in a strand-specific manner, that cover each nucleotide within a user-defined window (default: 100nt into the putative intronic region (upstream of 5’ss and downstream of 3’ss) and 200nt into the putative exonic region (downstream of 5’ss and upstream of 3’ss). See below for the analytical determination of the optimal default window sizes defined for this study.

#### Step 2: Changepoint detection algorithm for predicting putative exon boundaries

Exon boundaries were predicted using an algorithm to detect changepoints in read coverage that would be caused when intronic regions are spliced out around an exon with relatively consistent boundaries. Specifically, the *cpt.mean* function with “method = BinSeg, Q = 2” from the *Changepoint*^35^ and *Segmented*^36^ R packages was run on the base-specific read counts obtained in Step 1. Under these parameters, this function generates only two changepoints within a read coverage region, and gives no result if at least two breakpoints cannot be identified.

#### Step 3: Match putative exons predicted from 3’ and 5’ splice sites

Step 2 results in independent predictions of putative exons independently predicted from 3’ and 5’ splice sites, which need to be combined to arrive at a consensus exon. To do so, *CRYPTID-exon* uses the 3’ss associated exons as a baseline and calculates the following metrics for all 5’ss within the same gene:

- *exon start deviations:* distance between the 3’ss predicted exon start and all exon starts predicted from 5’ss
- *exon end deviations:* distance between the 3’ss predicted exon end and all exon ends predicted from 5’ss
- *3’ splice site deviation:* distance between the 3’ss and the predicted exon start (putative 3’ss)
- *5’ splice site deviation:* distance between the 5’ss and the predicted exon end (putative 5’ss)

All pairwise exon predictions from 3’ and 5’ splice sites for which all 4 of these metrics were less than 20nt were retained for further analysis. If a 3’ss predicted exon could be matched with multiple 5’ss, multiple criteria were prioritized to determine the most likely match (in the order listed below):

1. a 5’ss predicted exon with the same exon boundaries as that predicted from the 3’ss (*exon start deviation* = 0 & *exon end deviation* = 0)
2. a 5’ss predicted exon with at least one matching exon boundary as that predicted from the 3’ ss (*exon start deviation* = 0 | *exon end deviation* = 0)
3. a 5’ss predicted exon with the least combined distance from exon boundaries predicted from the 3ss (min(|*exon start deviation*| + |*exon end deviation*|)
4. a 5’ss predicted exon with whose predicted exon boundaries match the 3’ and 5’ splice sites used for prediction (*3’ss deviation* = 0 & *5’ss deviation* = 0)
5. a 5’ predicted exon with at least one predicted exon boundary matching either the 3’ or 5’ splice site used for prediction (*3’ss deviation* = 0 | *5’ss deviation* = 0)
6. A 5’ predicted exon with the least combined distance between predicted exon boundaries at the 3’ and 5’ splice sites used for prediction (min(|*3’ss deviation*| + |*5’ss deviation*|)

#### Step 4: Determine confidence of exon boundaries

After matching 3’ and 5’ss predicted exons, *CRYPTID-exon* categorizes the resulting consensus exons as low, medium, or high based on the consistency between the four deviation metrics described above. High confidence exons are those with zero deviation for all four metrics. Medium confidence exons are those where (1) only one of the four deviation metrics has a non-zero value or (2) either exon boundary matches (exon start deviation = 0 & exon end deviation = 0) or splice site matches (3’ss deviation = 0 & 5’ss deviation = 0), regardless of the values for the other deviations. Low confidence exons are the remaining matched exons for which all four deviation metrics are less than 20nt.

### Applying *CRYPTID-exon* to detect exons in K562 and KNS60 cells

Canonical and cryptic splice sites were identified and characterized for nascent (4sU labeled and nuclear fraction), nuclear, mature (polyA-enriched), and cycloheximide-treated RNA from K562 and KNS60 cells using *CRYPTID-ss* as described previously.^2^ Only splice sites supported by at least two reads or detected in at least two replicates were used for *CRYPTID-exon* prediction and characterization. All K562 and KNS60 datasets were obtained from GSE302375 unless otherwise indicated. *CRYPTID-exon* predictions were then classified by whether they were associated with 2, 1, or 0 cryptic splice sites (where the last category of exons are associated with 2 canonical sites). Predicted exons were further classified by their overlap with annotated exons, by using bedtools intersect to overlap predicted exon coordinates with annotated exon coordinates from ENSEMBL GRCH38.95:

- *annotated*: predicted exons whose boundaries precisely match the boundaries of an annotated exon
- *partial*: predicted exons whose boundaries are within the boundaries of an annotated exon
- *extended*: predicted exons whose boundaries overlap an annotated exon but add on additional exonic region(s) either upstream, downstream, or on both ends of the annotated exon
- *novel*: predicted exons whose boundaries do not partially or fully overlap annotated exon boundaries

### Identification of canonical exons and selection of optimal coverage windows

Exons expressed in K562 and KNS60 cells were identified using stringtie v2.2.1^50^ guided by reference-guided transcript assembly performed with ENSEMBL GRCH38.95 annotations. Identified exons were matched with GRCH38.95 annotations to identify canonical exons expressed in the datasets used.

Canonical exons were divided into 3 length bins: (1) exons ≤100nt, (2) 100 > exons ≤200nt, and (3) 200 > exons ≤ 500nt. 200 exons were randomly selected from each category and, base-specific coverage was obtained for 8 windows surrounding both the start and end of exons: +/- 100, 150, 200, 250, 300, 350, 400, 450nt. Exon boundaries were predicted with *CRYPTID-exon* using each of these coverage ranges, and used to calculate the prediction error (difference between predicted exon ends and actual exon ends) for each range.

### Identification of exons from multi-junction reads

To identify exons supported by multi-junction reads, uniquely mapped junction reads containing more than one splice junction were isolated using the *jM:B:c* attribute with pysam v0.20.0,^51^ for which the attribute specified multiple intron motifs. Intron coordinates for splice junctions in each read were obtained using *jI:B:I* attribute and multi-junction derived exon coordinates were extrapolated using the coordinates of flanking introns.

### Matching CRYPTID-exon predictions to canonical or multi-junction supported exons

The 3’ss of each canonical or multi-junction derived exon, was used to identify *CRYPTID-exon* predicted exons associated with that 3’ss and evaluate whether the canonical or multi-junction derived exon and the predicted exon shared perfect or fuzzy matched (less than or equal to 20 nt absolute difference between respective exon boundaries) exon boundaries.

### Validation of CRYPTID-exon predictions using long-read RNA sequencing data

To validate the presence of novel exonic regions in full-length RNA transcripts, we used polyA-enriched ISO-seq Pacific Biosciences long-read data from K562 cells (ENCODE; accession numbers: ENCFF429VVB, ENCFF634YSN, ENCFF694INI, ENCFF696GDL, and ENCFF763VZC). For each gene with a novel exonic region (associated with either canonical or cryptic splice sites), reads that mapped to that gene were filtered for those that started upstream and ended downstream of the novel exonic region to avoid biases from 5’ or 3’ truncation common in long-read data.^52^ Reads whose sequence covered 50% of the novel exonic region were considered to include the novel exonic region. Percent Spliced In (PSI) values for each novel exonic region was calculated as the number of reads that included the exon divided by the total number of filtered reads for the gene. To calculate the overall proportion of novel exonic regions present by long-read data, only exonic regions supported by a minimum of two reads were considered.

### Conservation analysis

For conservation analyses, we used two metrics. First, conservation scores for cryptic exons were obtained by applying the *bigWigAverageOverBed* tool from *ucsctools* to extract the mean phastCons score for each exon region using a hg38 phastCons (100 mammal alignment) bigwig file downloaded from the UCSC database.^53,54^ Second, base-specific conservation scores were obtained by applying the *bigWigToBedGraph* tool from ucsctools to extract base-specific phyloP scores for the 20nt window surrounding exon starts (3’ splice sites) and exon ends (5’ splice sites) of exons using a hg38 phyloP (100 mammal alignment) bigwig file downloaded from the UCSC database.^53,54^

### Estimating percent inclusion of *CRYPTID-exon* predicted exons

To evaluate the extent to which cryptic exons were included in full-length transcripts, we used two versions of a percent spliced in (PSI) metric:

(1) PSI coverage (Ѱ_cov_): The first metric relies on estimating a PSI from read coverage in and flanking the cryptic exon. This is specifically designed for calculating PSI values from nascent or nuclear RNA populations, where there is a combination of spliced (excluding and including cryptic exons) and not yet spliced molecules. This method assumes that while reads within the exon represent both spliced and not-yet-spliced molecules, intronic regions around the cryptic exon are representative of only molecules that have not yet been spliced. Specifically, the method first calculates cryptic exon inclusion and exclusion values:

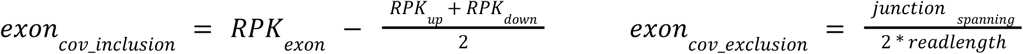

Where 𝑅𝑃𝐾_𝑒𝑥𝑜𝑛_ is the reads per kilobase within the cryptic exon, 𝑅𝑃𝐾_𝑢𝑝_ and 𝑅𝑃𝐾_𝑑𝑜𝑤𝑛_ are the reads per kilobase from the 100nt regions upstream and downstream of the cryptic exon (respectively), 𝑗𝑢𝑛𝑐𝑡𝑖𝑜𝑛_𝑠𝑝𝑎𝑛𝑛𝑖𝑛𝑔_ is the number of junction reads entirely spanning the exon without intersecting it, and 𝑟𝑒𝑎𝑑𝑙𝑒𝑛𝑔𝑡ℎ is the length of the reads. All read counts were obtained using bedtools coverage ^49^ with the “-split” parameter (except in the case of the junction reads). The PSI value was then calculated as follows:

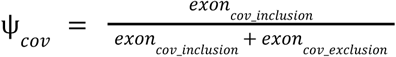

(2) PSI junction (Ѱ_junc_): The second metric relies on estimating a PSI using only junction reads supporting the inclusion or exclusion of the cryptic exon, explicitly excluding the contribution from any molecules that have not yet been spliced. Specifically, the method first calculates cryptic exon inclusion and exclusion values:

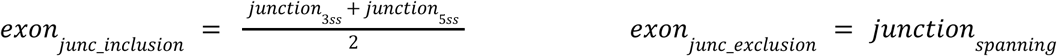

Where 𝑗𝑢𝑛𝑐𝑡𝑖𝑜𝑛_3𝑠𝑠_ and 𝑗𝑢𝑛𝑐𝑡𝑖𝑜𝑛_5𝑠𝑠_ are the number of junction reads supporting the 3’ and 5’ splice sites (respectively) and 𝑗𝑢𝑛𝑐𝑡𝑖𝑜𝑛𝑠_𝑠𝑝𝑎𝑛𝑛𝑖𝑛𝑔_ is the number of junction reads entirely spanning the exon without intersecting it. Reads supporting the 3’ and 5’ splice sites were obtained from the output of *CRYPTID-ss*, while spanning junction read counts were obtained as described above. The PSI value was then calculated as follows:

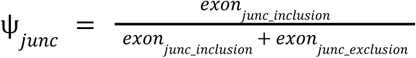

### Evaluating translational reading frames of cryptic exons

The sequence for each *CRYPTID-exon* prediction in a protein-coding gene was obtained using bedtools getfasta^49^ with the “-s” parameter using hg38 reference genome. For each exon, the nearest upstream coding exon was identified using the isoform with the highest expression level (estimated with Kallisto 0.45.0^55^) in mature RNA. The frame of that upstream coding exon was then imposed upon the cryptic exon to screen for the presence of UAA, UAG, or UGA stop codons.

### Human cell culture

K562 cells were grown in IMDM supplemented with 10% FBS. KNS60 cells were grown in DMEM supplemented with 5% FBS. All cell lines were maintained at 37°C and 5% CO_2_ until 70% confluent.

### Amplicon sequencing to validate cryptic exons

Four primer sets were designed for each cryptic exon being validated using Primer3^56–58^ such that successful amplification would only occur in the presence of the cryptic exon (**Supplementary Table 2**). Total RNA was extracted as described previously^2^ and PCR amplification was performed using Phire Hot Start II PCR Master Mix (Thermofisher, F126) with a 55°C annealing temperature, 15second extension time, and 40 cycles. Completed reactions were run on 1% agarose gel. Selected amplicons were sent to Genewiz from Azenta Life Sciences for Amplicon-EZ Next-Generation Sequencing and sequenced on a MiSeq platform in 500-cycle paired-end format. Resulting sequences were then mapped to the hg38 reference genome with ENSEMBL GRCH38.95 reference annotations using STAR^59^ with default parameters and cryptic exon usage was confirmed using visualization of read coverage and junction reads across the region.

### Targeted sequencing of nuclear RNA from specific genes

Gene-specific transcript enrichment was performed following an adapted version of a previously published protocol.^60^ Specifically, RNA probes were used to enrich for target genes from cDNA from nuclear RNA and library preparation and sequencing was conducted on enriched samples, as detailed below.

#### Design of RNA probes for targeted enrichment

Regions of high conservation were identified within target gene sequences using the UCSC Genome Browser (https://genome.ucsc.edu/), ensuring consistent and effective target engagement across isoforms. The PFizer RNAi Enumeration and Design tool (PFRED, https://github.com/pfred) was used to generate 20 antisense sequences with a high probability of effective RNA engagement based on secondary structure and various thermodynamic properties (**Supplementary Table 3**). This tool is designed to identify effective siRNA/ASO sequences for target degradation, but we expect that similar properties also enable effective binding to target transcripts for pulldown. RNA probes were synthesized in-house, using 40-mers sequences to encourage highly specific binding, while minimizing intramolecular secondary structure. A biotin moiety separated by a hexaethylene glycol (HEG) molecule to encourage mobility was added to the 3’ end of the RNA probe to facilitate streptavidin-based enrichment.

#### Reverse transcription and target enrichment

Reverse transcription of nuclear RNA from KNS60 cells isolated as described in Khokhar *et al.*^2^ was performed using the High-Capacity cDNA Reverse Transcription Kit (Thermofisher, 4368814) according to manufacturer specifications. Single-stranded cDNA was treated with alkaline hydrolysis buffer (150mM NaOH, 1mM EDTA) to degrade any remaining RNA template. 10X TE buffer, pH 7.4 treatment for 10 minute incubation at 50°C was used to neutralize the hydrolysis reaction. cDNA was purified and buffer-exchanged into 1X TE buffer using 3K MWCO Amicon cartridges. The purified cDNA was added to 1 Unit of SUPERase IN RNase Inhibitor per 100ng cDNA (Invitrogen, AM2696), 1pmol biotinylated probe mix per 16ng cDNA, and 200uL per 400ng cDNA of hybridization/wash buffer (40mM Tris-HCl, 300mM NaCl, 0.2% Tween 20 (w/v), 1% SDS) and the sample was incubated at 60°C for 20 minutes with regular agitation. 500ug of magnetic streptavidin beads (Pierce, 88816) per 400ng of cDNA were added to the reaction and the reaction was again incubated at 60°C for 20 minutes with regular agitation. Following incubation, the beads were immobilized and washed 3x with hybridization/wash buffer. Alkaline hydrolysis buffer was added to destroy the RNA probes and release the enriched cDNA transcripts. 10X TE buffer was used to neutralize the hydrolysis reaction and samples were buffer exchanged into 1X TE using 3K MWCO Amicon cartridges. Samples were purified using ethanol precipitation (2.67X volumes of 100% EtOH and 0.1X volumes 3M sodium acetate).

#### Second-strand synthesis and library preparation

DNA Polymerase I (*E. coli*) was used to generate second-strand cDNA from the enriched, single-stranded cDNA. The following reagents were combined: 10X NEBuffer 2, 25mM dNTPs, 10X random primers, DNA polymerase I (NEB, M0209), and water for a total reaction volume of 50uL. The reaction was incubated at 95°C for 60 seconds, 16°C for 4 hours, and 37°C for 1 hour. Second-strand cDNA was ethanol precipitated (2.67X volumes of 100% EtOH and 0.1X volumes 3M sodium acetate) and library preparation was performed using NEBNext Ultra II FS Library Prep Kit. Libraries were sequenced on a NextSeq2000 using 2×100 reads.

#### Computational analysis of target enriched nuclear RNA-seq

Samples were analyzed as described in Khokhar et al.^2^ Briefly, samples were PEAR-procssed,^61^ subsampled to 112M single reads with seqtk^62^, mapped using STAR with parameters specified in Khokhar et al.^2^ Kallisto^55^ was used to estimate gene expression levels in the target-enriched samples and confirm enrichment relative to whole nuclear RNA. Samples were analyzed with *CRYPTID-ss* to obtain high-confidence cryptic splice sites and processed with *CRYPTID-exon* to predict exons associated with the detected cryptic splice sites. Exons from the sites with the highest usage were chosen for downstream experimentation following visual inspection of the region and validation by amplicon sequencing as described above.

### Antisense oligonucleotide (ASO) treatments

#### ASO design

ASOs were designed to induce skipping of the targeted cryptic exon for *NF1* and *TBK1* using 3 different methods: (1) eSkip-Finder, from which the top 10 ASO sequences per gene with the highest predicted efficacy were selected,^40^ (2) ESEfinder 3.0 was used to identify putative exon splicing enhancers (ESEs) and 5 ASOs per gene were designed to occlude these ESEs,^41,63^ and (3) custom ASOs were designed to occlude either the 5’ end of the cryptic exon or the predicted 5’ or 3’ splice sites. In total, 20 ASOs were designed for each cryptic exon (**Supplemental Table 4**). All ASOs were synthesized in-house with a phosphorothioate (PS) backbone and 2’-O-methoxyethyl (2’-MOE) ribose modifications.

#### ASO synthesis

Oligonucleotides were synthesized using phosphoramidite chemistry on a DrOligo 48 synthesizer (Biolytic, Fremont, CA) using standard methods.^64^ All oligonucleotides were synthesized at a 1 𝜇mole synthesis scale using 1000 Å Unylinker-functionalized long-chain alkyl amine (LCAA) controlled pore glass (CPG) solid supports (ChemGenes). Phosphoramidites were purchased from ChemGenes (Wilmington, MA) and diluted to 0.1 M in anhydrous acetonitrile (ACN). A solution of 0.25 M 5-(benzylthio)-1H-tetrazole (BTT) in acetonitrile was used as the activator, and the coupling time for all phosphoramidites was 4 minutes. Detritylation was achieved using 3% trichloroacetic acid in dichloromethane. Capping was achieved using 20% n-methylimidazole in ACN (Cap A), and 20% acetic anhydride and 30% 2,6- lutidine in ACN (Cap B). Reagents for capping and detritylation were purchased from ChemGenes. Oxidation was performed using 0.05 M iodine in pyridine-H_2_O (9:1, v/v), or sulfurization with 0.1 M [(Dimethylamino-methylidene)amino]-3H-1,2,4- dithiazoline-3-thione (DDTT) in pyridine (both ChemGenes).

#### ASO deprotection, desalting, quantification, and quality control

Backbone cyanoethyl deprotection was achieved on the support by treatment with 10% diethylamine in ACN. Oligonucleotides were cleaved from the solid support and base-protecting groups removed by treatment with conc. aqueous NH_4_OH (Sigma-Aldrich) at 55°C for 18 hours. Oligonucleotides were dried using centrifugal evaporation, resuspended in water, and transferred to 3kDa molecular weight cut-off Amicon filters (Millipore) for desalting. Oligonucleotides were washed once with PBS and twice with water, then transferred to microcentrifuge tubes and stored at -20°C until use.

Oligonucleotides were quantified by measuring absorbance at 260 nm using a Nanodrop One (ThermoFisher). Concentrations were calculated using the Beer-Lambert equation using extinction coefficients calculated for each sequence using published values. The identities of oligonucleotides were confirmed by LC–MS analysis using an Agilent 6530 accurate mass Q-TOF using the following conditions: buffer A: [100 mM 1,1,1,3,3,3-hexafluoroisopropanol (HFIP) and 9 mM triethylamine (TEA) in LC–MS grade water]; buffer B: [100 mM HFIP and 9 mM TEA in LC–MS grade methanol] using a linear gradient of 5–100% B over 5 minutes at 60°C; column, Agilent AdvanceBio oligonucleotides C18; flow rate, 0.85 ml/min. LC peaks were monitored at 260 nm. MS parameters: Source, electrospray ionization; ion polarity, negative mode; range, 100–3200 m/z; scan rate, 2 spectra/s; capillary voltage, 4000 V; fragmentor, 200 V; gas temp, 325°C.

#### ASO treatment

Single ASO transfections in KNS60 cells were performed with Lipofectamine RNAiMAX (Invitrogen, 13778150) with ASOs against their respective targets at a final concentration of 80nM or 160nM (from an ASO stock concentration of 20uM or 40uM). The stock ASO concentration:lipid ratio was 4:3. Reagents were combined in Opti-MEM and introduced to cells 24 hours after plating and incubated at 37°C and 5% CO_2_ for 24 or 48 hours. When cells were treated with 2 ASOs simultaneously, the ASOs had a combined concentration of either 80nM or 160nM. Transfections were also performed with a scrambled ASO (non-targeting control) and a lipid-only control. A non-transfected control (untreated) was included as an additional comparator group.

#### Analysis of ASO effects

Changes in gene expression levels of target genes were estimated using Taqman RT-qPCR with predesigned PrimeTime qPCR probe assays (IDT) for *NF1* (Hs.PT.58.27908857) and *TBK1* (Hs.PT.58.25458154), using total RNA from transfected cells (Supplementary Table 5). *HPRT* (Hs.PT.58v.45621572) was used as a housekeeping control gene for normalization. To characterize differential splicing of cryptic exons following ASO-treatment at 80nM for 24 hours, nuclear RNA from ASO-treated and untreated control KNS60 cells was used to selectively enrich target genes, followed by second-strand synthesis as described above. The non-targeting control ASO group was used for final normalization. Libraries were prepared using the NEBNext Ultra II FS DNA Library Prep Kit for Illumina, inputs <100ng (NEB, E7805) according to manufacturer’s instructions. Libraries were sequenced as 2×100 reads on a NextSeq 2000 using NextSeq 1000/2000 P2 Reagents (200 cycles) v3 (Illumina, 20046812).

### Computational analysis of ASO sequencing data

Samples were analyzed as described in Khokhar *et al.*^2^ Briefly, samples were PEAR-procssed^61^, subsampled to 10.7M single reads with seqtk^62^, and mapped using STAR with parameters specified in Khokhar *et al.*^2^ Cryptic splice sites were identified *CRYPTID-SS* as described in Khokhar *et al.*^2^ and cryptic exons were identified with *CRYPTID-exon*, using a maximum deviation value of 100 to cast a wider net for potential cryptic exons.

## CODE & DATA AVAILIBILITY

The *CRYPTID*-exon pipeline is available at https://github.com/thepailab/CRYPTID-exon. Sequencing data generated for this study are available at the Gene Expression Omnibus (GSE312830).

## ACKNOWLEDGEMENTS

We thank Allan Jacobson and members of the Watts and Pai labs for helpful discussions and comments. This work was funded by grants from the National Institutes of Health (R35GM133762 A.A.P. and R01NS111990 to J.K.W.), and the National Science Foundation (CAREER2237568 to A.A.P.).

## AUTHOR CONTRIBUTIONS

Contributed equally (ESK, KB), Conceived and designed experiments (JKW, AAP), Performed the experiments (KB, ZK, AH, JL, AW), Performed statistical analyses (ESK, ECR, AAP), Analyzed the data (ESK, KB, MK, ECR, AAP), Contributed reagents/materials/analysis tools (JKW, AAP), Wrote the paper (ESK, JKW, AAP), Jointly supervised research (JKW, AAP).

## Notes

### Competing Interest Statement

The authors have declared no competing interest.

