## Supplementary Material for "*CRYPTID-exon*: streamlined detection of cryptic exons from RNA-seq data"

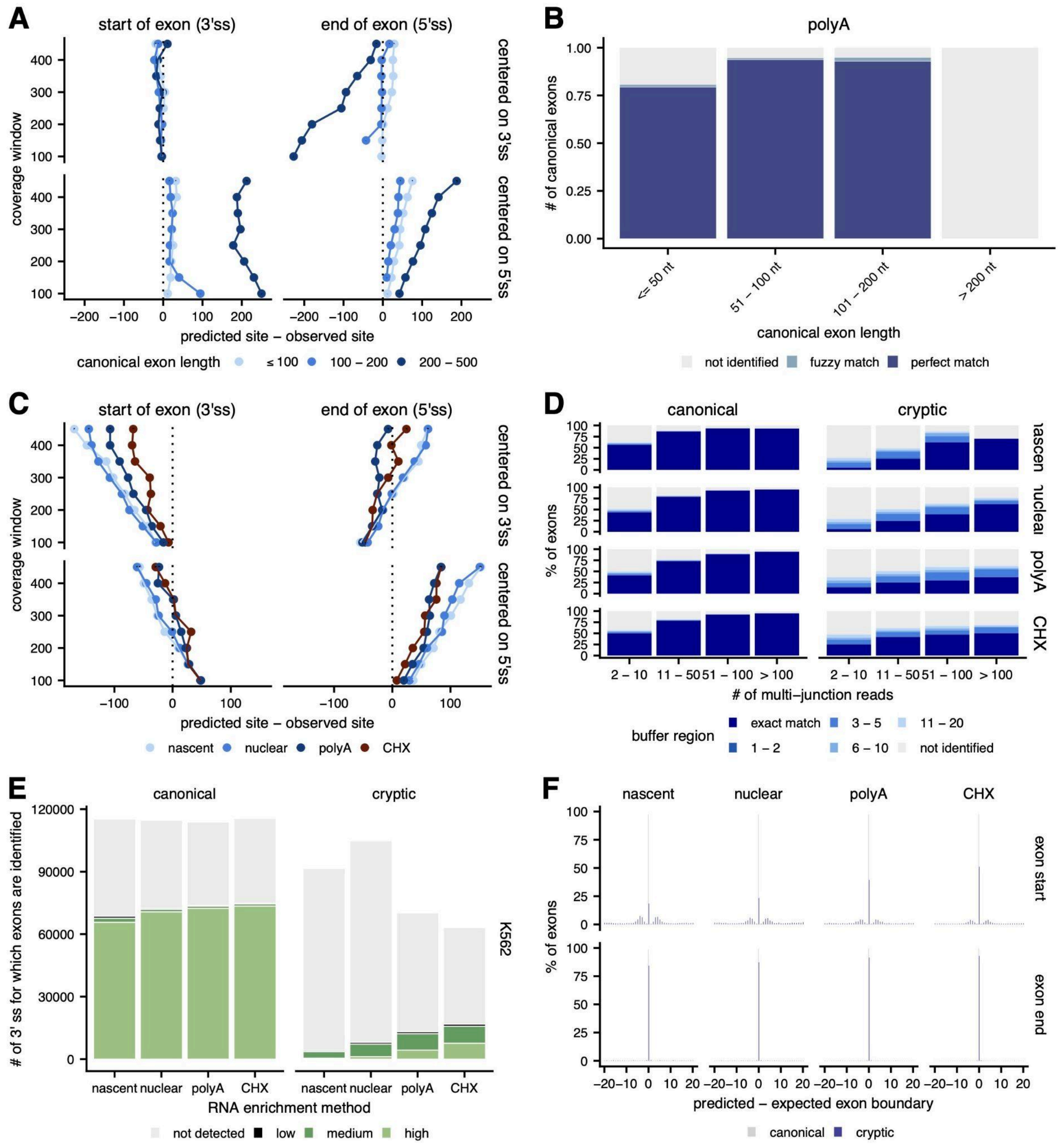


**Figure S1. *CRYPTID-exon* identifies exon boundaries with high specificity. (A)** Difference between the predicted exon boundary and the empirically observed canonical splice site (*x-axis*) across different coverage windows used for *CRYPTID-exon* identification (*y-axis*). For each bin of canonical exon lengths (*shades of blue*), 200 exons were randomly sampled among all canonical exons expressed in K562 mature RNA. **(B)** Percentage of canonical exons correctly identified by *CRYPTID-exon*, with either perfect (*dark blue*) or fuzzy boundary (*light blue*) matches within each category of exon length (*x-axis*) across RNA enrichment methods. Canonical exons are those expressed in K562 mature RNA. **(C)** Difference between the predicted exon boundary and the empirically observed cryptic splice site (*x-axis*) across different coverage windows used for *CRYPTID-exon* identification (y-axis) in 200 exons randomly sampled from multi-junction derived exons across RNA enrichment methods. **(D)** Percentage of canonical or cryptic exons from multi-junction reads correctly identified by *CRYPTID-exon*, separated by buffer region distance (*shades of blue*), across ranges of supporting multi-junction reads and RNA enrichment methods. **(E)** Number of canonical (*left*) or cryptic (*right*) exons identified by *CRYPTID-exon*, separated into different confidence levels (*shades of green*) across RNA enrichment methods from K562 and KNS60 cells. **(F)** The distribution of the difference between the *CRYPTID-exon* predicted exon boundary and the expected exon boundary (from empirically observed splice site) for canonical (*grey*) or cryptic (*blue*) exons across RNA enrichment methods.

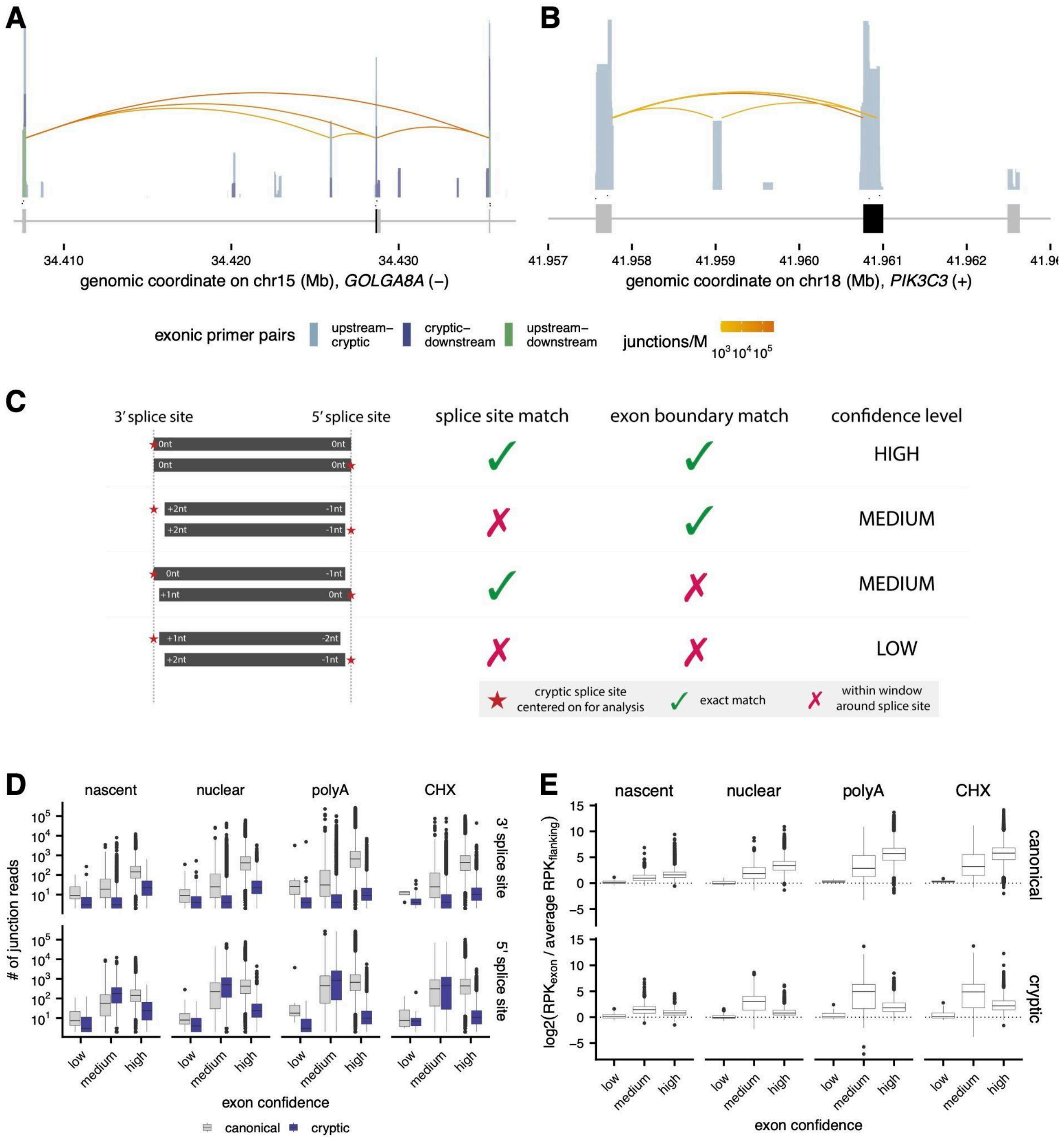


**Figure S2**. ***CRYPTID-exon* identifies exon boundaries across confidence levels. (A, B)** Amplicon-seq read coverage (*top*) validating cryptic exons identified in *GOLGA8A* **(A)** and *PIK3C3* **(B)** in K562 cells, with splice junction reads (*curved lines*, shaded by abundance), primer pairs used for amplication (*middle arrows*), and gene annotations (*bottom*). Coverage is colored by the positioning of the primer pairs relative to the cryptic exon. **(C)** Schematic showing how low, medium, and high confidence exons are categorized. **(D)** Number of junction reads supporting empirically observed splice sites used to identify canonical (*grey*) or cryptic (*blue*) exons across exon confidence levels and RNA enrichment methods. **(E)** The fold change of normalized exonic coverage relative to flanking intronic (average of 100nt upstream and downstream of predicted exon) across exon confidence levels and RNA enrichment methods.

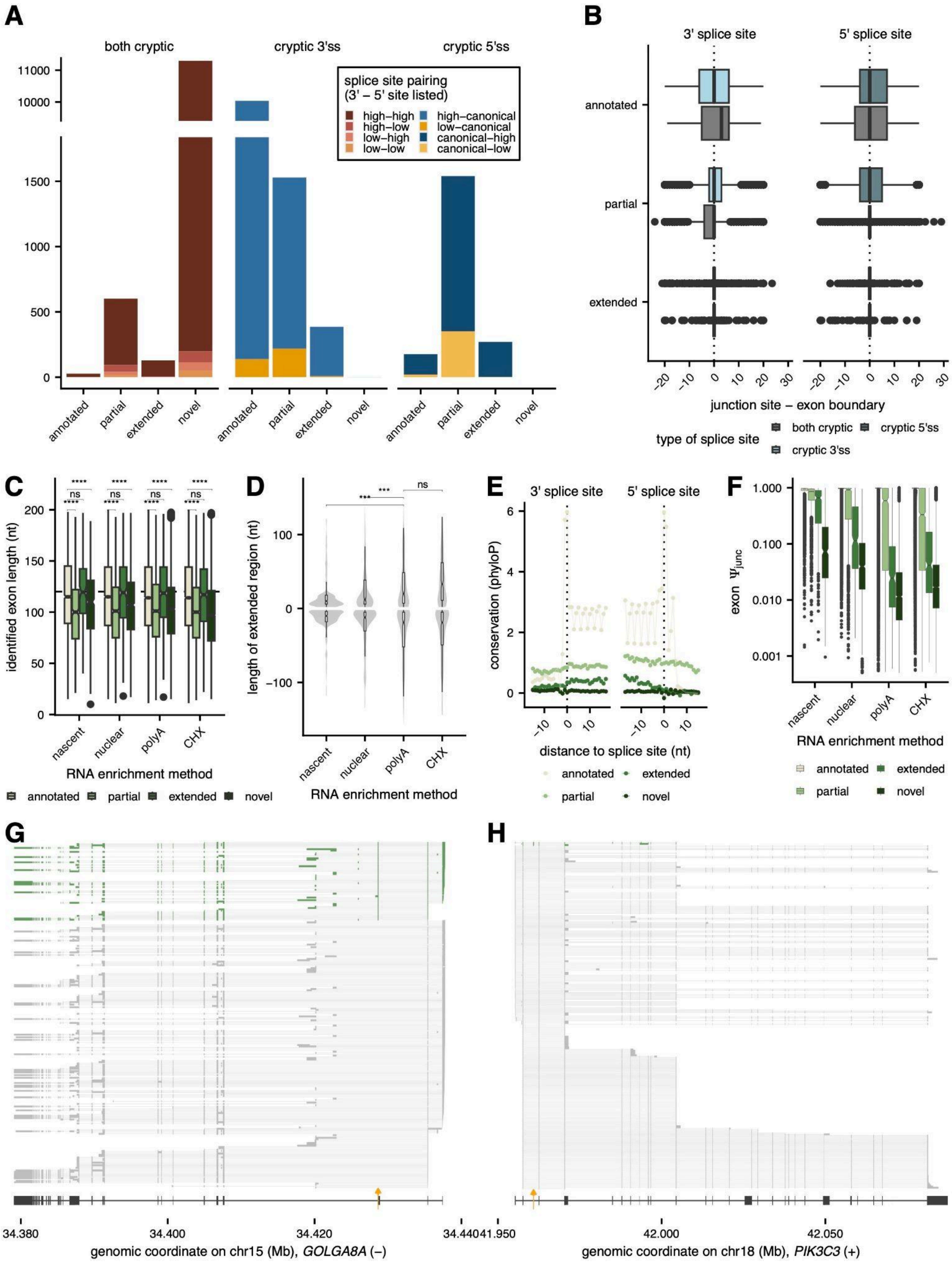


**Figure S3.** ***CRYPTID-exon*** **identifies novel exonic regions associated with cryptic splice sites across RNA enrichment methods. (A)** Number of cryptic exons within each overlap category for exons with two cryptic splice sites (*left*), a cryptic 3'ss (*middle*), or cryptic 5'ss (*right*), with colors indicating the fidelity and pairing of splice sites across the exon. **(B)** Distance between empirically observed splice sites and predicted exon boundaries across overlap categories. **(C)** Distribution of cryptic exon lengths across overlap categories (*shades of green*) and RNA enrichment methods. **(D)** Distribution of the extended region lengths for cryptic exons that use alternative 3' or 5' splice sites relative to a canonical exons across RNA enrichment methods. **(E)** Nucleotide-specific phyloP scores (*y-axis*) in a 20nt window around 5′ and 3′ splice sites across cryptic exon overlap categories (*shades of green*). **(F)** Distribution of exon $\Psi_{junc}$ values for cryptic exons across overlap categories (*shades of green*) and RNA enrichment methods. **(G, H)** Iso-Seq long-read sequencing reads for *GOLGA8A* **(G)** and *PIK3C3* **(H)**, with gene annotations shown at the bottom and the cryptic exon position marked by the yellow arrow. Reads including the cryptic exon are in green. Signficance was assessed with a Mann-Whitney U Test. *** adjusted p-value < 0.001, **** adjusted p-value < 0.0001.

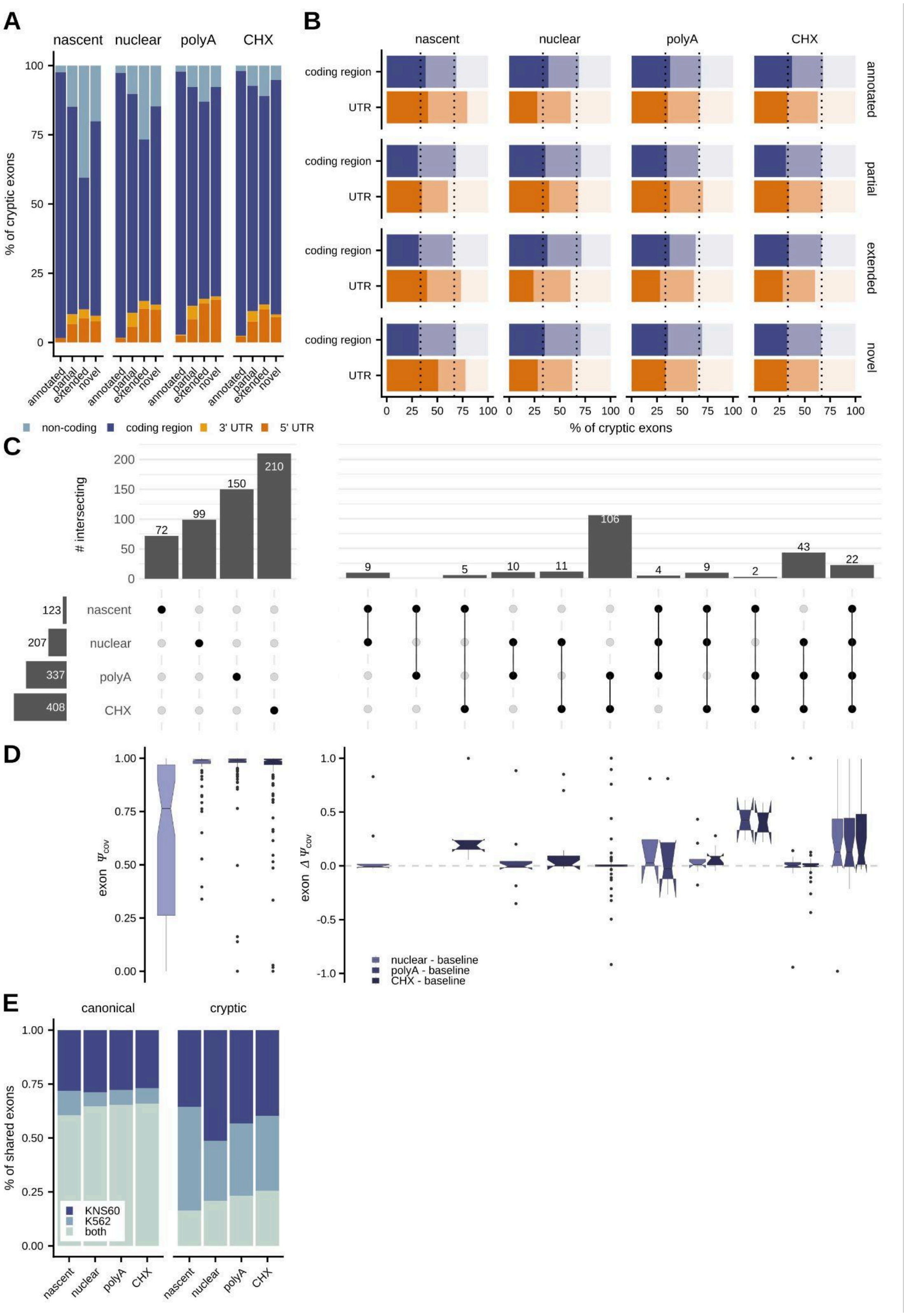


**Figure S4. RNA enrichment method and cell-type-specific usage of cryptic exons. (A)** Percentage of cryptic exons that are found in non-coding genes (*light blue*), coding regions (*dark blue*), 3'UTRs (*light orange*), or 5' UTRs (*dark orange*) across overlap categories and RNA enrichment methods. **(B)**

Percentage of cryptic exons in coding regions (*blue*) or UTRs (*orange*) whose length is evenly divisible by 3 (mod3 = 0; maintaining frame; *darkest shade*) or not evenly divisible by 3 (mod3 = 1 or mod3 = 2; *lighter shades*) across overlap categories and RNA enrichment methods. **(C)** The number of extended cryptic exons detected in only one (*left*) or multiple (*right*) RNA enrichment methods, with the total number of extended cryptic exons within each RNA enrichment method shown as bars on the left. **(D)** Distribution of exon $\Psi_{cov}$ values (*left*) or exon $\Delta\Psi_{cov}$ values (*right*) for extended cryptic exons that show different overlap patterns (as shown in C). Exon $\Delta\Psi_{cov}$ values are estimated relative to the earliest RNA fraction in which the exon is identified. **(E)** Proportion of cryptic exons identified only in K562 (*light blue*), only in KNS60 (*dark blue*), or shared between both cell types (*green*).

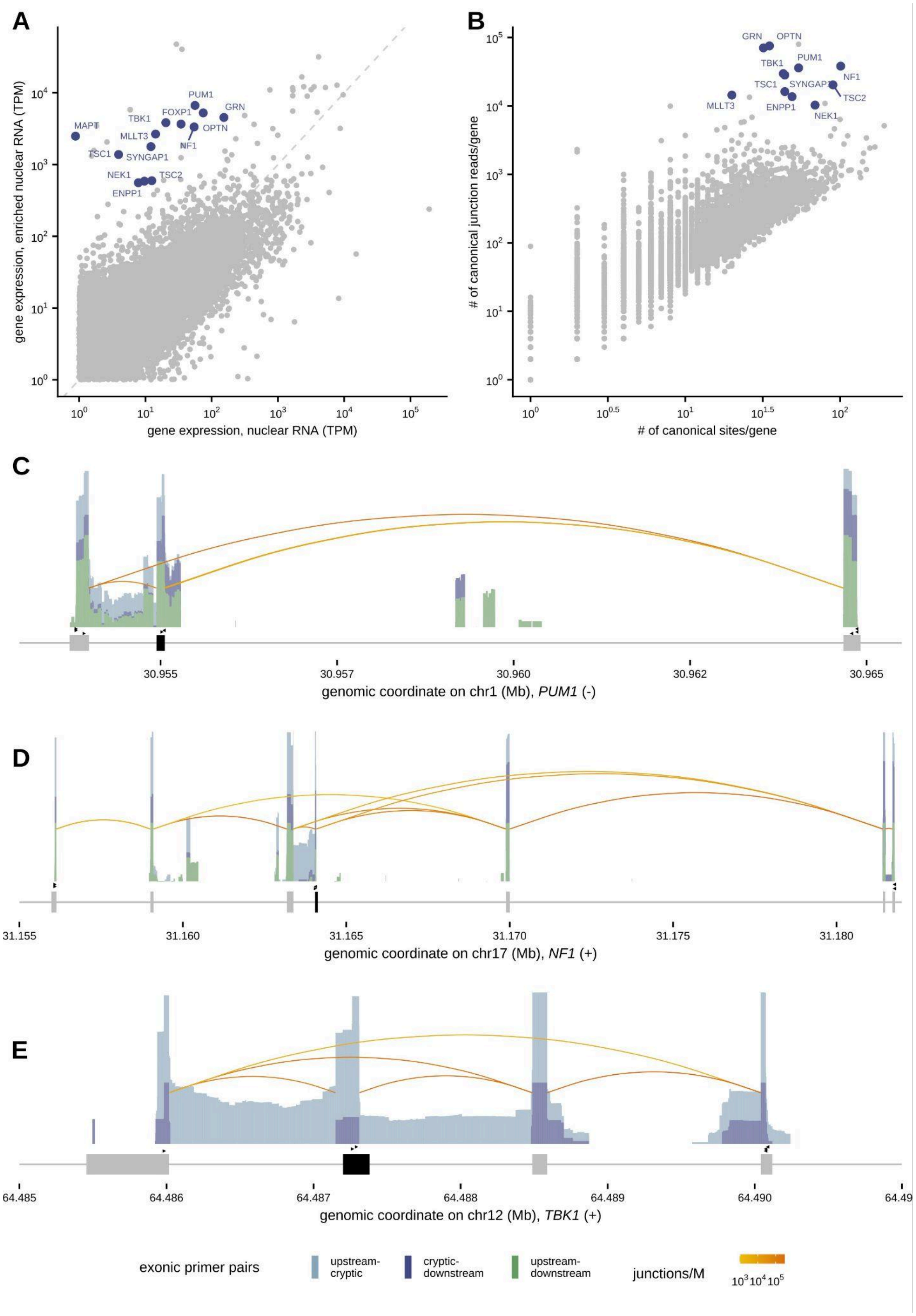


**Figure S5. Validation of cryptic exons identified through targeted sequencing of disease-relevant genes. (A)** Gene expression in nuclear RNA (*x-axis*) vs enriched nuclear RNA (*y-axis*), with target genes highlighted in blue. **(B)** The number of canonical sites identified per gene (*x-axis*) vs. the number of canonical junction reads identified per gene (*y-axis*) in enriched nuclear RNA, with target genes highlighted in blue. **(C, D, E)** Amplicon-seq read coverage validating cryptic exons identified in *PUM1* **(C),** *NF1* **(D)**, and *TBK1* **(E)** in KNS60 cells, with splice junction reads (*curved lines*, shaded by abundance), primer pairs used for amplification (*middle arrows*), and gene annotations (*bottom*). Coverage is colored by the positioning of primer pairs relative to the cryptic exon

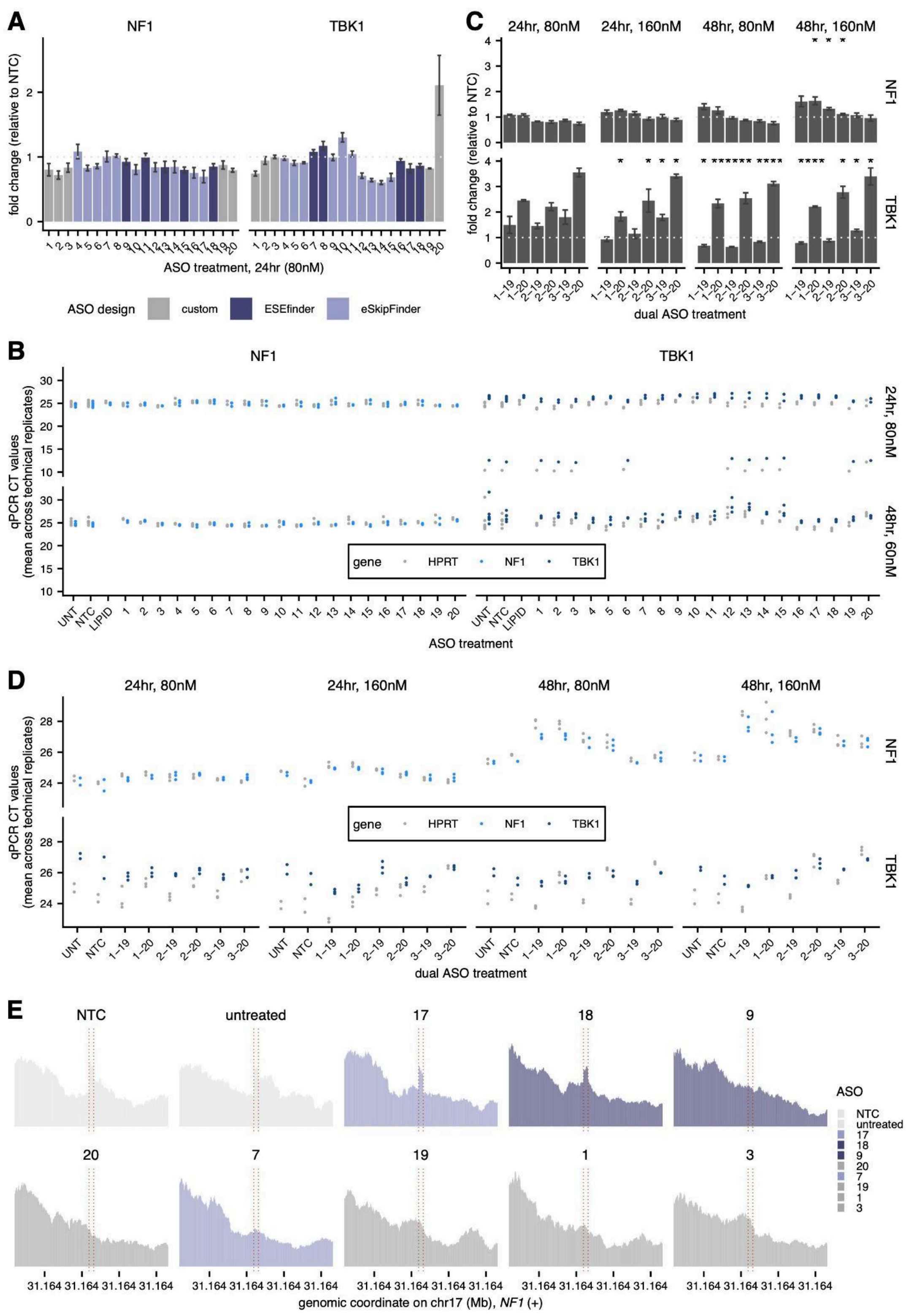


**Figure S6. Gene expression and splicing changes in *NF1* and *TBK1* following treatment with ASOs. (A)** Fold changes in gene expression levels estimated by RT-qPCR for *NF1* (*left*) and *TBK1* (*right*) relative to a non-treated control (NTC), following transfection with individual ASOs at 80nM for 24 hours. ASOs are numbered by their genomic position, with 1 and 20 representing the upstream-most and downstream-most ASOs, respectively. **(B)** CT values (*y-axis*) from quantitative RT-PCR after treatment with individual ASOs for conditions specified in A. **(C)** Fold changes in gene expression levels estimated by RT-qPCR for *NF1* (*top*) and *TBK1* (*bottom*) relative to NTC, following joint transfection with two ASOs at 80nM for 24 hours (*left*), 160nM for 24 hours (*left middle*), 80nM for 48 hours (*right middle*), and 160nM for 48 hours (*right*). **(D)** CT values (*y-axis*) from quantitative RT-PCR after joint transfection with two ASOs for conditions specified in B. **(E)** Read coverage around NF1-identified cryptic exon (*y-axis*) in enriched nuclear RNA following ASO treatment or in control samples. Significance was assessed with a paired T-test. * FDR < 0.1, ** FDR < 0.01, *** FDR < 0.001.
